# Stochastic Modeling Reveals Kinetic Heterogeneity in Post-replication DNA Methylation

**DOI:** 10.1101/677013

**Authors:** Luis Busto-Moner, Julien Morival, Arjang Fahim, Zachary Reitz, Timothy L. Downing, Elizabeth L. Read

**Affiliations:** Institut Quimic de Sarrià, Universitat Ramon Llull, Barcelona, Spain; Dept. of Chemical & Biomolecular Engineering, University of California, Irvine, California, USA; Dept. of Biomedical Engineering, University of California, Irvine, California, USA; Center for Complex Biological Systems, University of California, Irvine, California, USA; NSF-Simons Center for Multiscale Cell Fate Research, University of California, Irvine, California, USA

## Abstract

DNA methylation is a heritable epigenetic modification that plays an essential role in mammalian development. Genomic methylation patterns are dynamically maintained, with DNA methyltransferases mediating inheritance of methyl marks onto nascent DNA over cycles of replication. A recently developed experimental technique employing immunoprecipitation of bromodeoxyuridine labeled nascent DNA followed by bisulfite sequencing (Repli-BS) measures post-replication temporal evolution of cytosine methylation, thus enabling genome-wide monitoring of methylation maintenance. In this work, we combine statistical analysis and stochastic mathematical modeling to analyze Repli-BS data from human embryonic stem cells. We estimate site-specific kinetic rate constants for the restoration of methyl marks on >10 million uniquely mapped cytosines within the CpG (cytosine-phosphate-guanine) dinucleotide context across the genome using Maximum Likelihood Estimation. We find that post-replication remethylation rate constants span approximately two orders of magnitude, with half-lives of per-site recovery of steady-state methylation levels ranging from shorter than ten minutes to five hours and longer. Furthermore, we find that kinetic constants of maintenance methylation are correlated among neighboring CpG sites. Stochastic mathematical modeling provides insight to the biological mechanisms underlying the inference results, suggesting that enzyme processivity and/or collaboration can produce the observed kinetic correlations. Our combined statistical/mathematical modeling approach expands the utility of genomic datasets and disentangles heterogeneity in methylation patterns arising from replication-associated temporal dynamics versus stable cell-to-cell differences.

## Introduction

DNA methylation is an essential epigenetic modification found in a diversity of organisms, which is broadly associated with silencing of genes [1]. Methylation patterns across the genome encode epigenetic information associated to cellular processes including differentiation [2, 3] and genomic imprinting [4, 5]. These patterns are also conserved in distinct cell types, and clearly distinguish cell types in mammalian tissues [6–8]. Failure in the transmission of such patterns from one generation to the next and the appearance of aberrant methylation patterns have been associated with cancer [9, 10], aging [11], or organismal death [12].

In mammals, DNA methylation is primarily found in the cytosine-phosphate-guanine (CpG) dinucleotide context, which presents a symmetric substrate for inheritance that echoes the Watson-Crick model of genetic inheritance [12, 13]. Methylation patterns are generally transmitted with high fidelity from the parent template strand to nascent DNA over cycles of DNA replication. The classic model of *maintenance* methylation holds that DNA Methyltransferase 1 (DNMT1) is primarily responsible for this inheritance, which it accomplishes by localizing to replication foci [14] and preferentially catalyzing addition of methyl groups onto hemi-methylated CpG substrates (i.e., those CpG substrates with methylation present on only the parent-strand cytosine) [15–17]. In contrast to DNMT1, DNMT3A and 3B are often termed *de novo* methyltransferases because their catalytic activity shows no preference for hemimethylated versus unmethylated DNA and they are essential in the establishment of genome-wide methylation patterns during embryogenesis [18]. However, in recent years it has been pointed out that this classical model is overly-simplistic [19], since, for example, DNMT1 and DNMT3s are both essential for development, both contribute to maintenance methylation [20], and these enzymes work together with methyl-eraser enzymes (Ten-eleven translocation proteins (TETs) [21]) to control methylation across the genome and over time.

Whole genome bisulfite sequencing, which maps the methylation status of individual CpGs, shows generally bimodal patterns comprising fully methylated or fully demethylated regions. That is, the fraction of cells in a population with methylation at a given site tends to be near 1 or 0. However, intermediate methylation (IM), where methylation fraction is between 0 and 1, is also widespread. Despite broad conservation of genomic methylation patterns in distinct cell types, some loci show this type of non-uniformity in methylation across homogeneous cell populations. This heterogeneity appears to be itself conserved, as common IM regions have been identified across individuals and even species [22]. IM appears to be critical for proper organism development and cell fate determination [22–24], contributes to genomic imprinting [8, 25], and plays a prominent role in tumor cell evolution [26].

The determinants of IM are not fully understood. In some contexts, cell-to-cell heterogeneity within populations has been implicated as the chief contributor to IM [27, 28]. However, as methylation levels result from dynamic processes carried out asynchronously in different cells, IM could result not only from stable cell-to-cell differences, but also from temporal heterogeneity. For example, in an unsynchronized population of replicating cells, a subset of cells would be in the process of re-establishing methylation marks post-replication, thus contributing to lowered methylation fractions at the bulk level. A recently-developed experimental technique, Replication-associated Bisulfite Sequencing (Repli-BS), enables time-resolved measurement of genomic methylation patterns, including in newly replicated DNA [29], shedding light on dynamic re-establishment of methylation that must occur after each round of DNA replication. Using this technique, Charlton and Downing, et al. reported a pronounced genome-wide delay of several hours in post-replication nascent strand DNA methylation in human Embryonic Stem Cells (hESCs). These results echoed previous observations of a lag in maintenance methylation following replication in a variety of mammalian cell types [20, 30–32]. Furthermore, Charlton and Downing, et al., reported that the delay in post-replication nascent strand methylation accounts for a significant amount of the IM observed in hESCs in WGBS experiments.

Along with experimental evidence, mathematical modeling has informed understanding of DNA methylation dynamics. Population epigenetic models have explored the interplay between processes including enzyme-mediated *de novo* methylation, maintenance methylation, demethylation, and replication [33–38]. Some models have incorporated various mechanisms of interdependence of CpGs, where, for example, the efficiency of maintenance methylation at a given site depends on the methylation status of its neighbors [39–44]. Biochemical studies have enabled the development of enzyme-kinetic models and parameter quantification for methyltransferase activity [17, 45, 46]. While a number of modeling studies based on *in vivo* data in various cellular contexts have quantified the relative efficiency of maintenance methylation (i.e., the probability that the methylated state is successfully propagated through one cell division cycle), genome-wide quantification of sub-cell-cycle kinetics of maintenance methylation *in vivo* has not been possible.

The expansion in recent years of genomic measurement techniques provides an increasingly fine-grained view of methylation patterns across the genomic, across cells, and across time. However, there remains a major gap in our understanding of the molecular sources and regulatory consequences of most of the heterogeneity present within the mammalian methylome. In this work, we combine statistical inference and mathematical modeling to analyze genome-wide post-replication methylation kinetics, making use of published Repli-BS data from hESCs. First, using Maximum Likelihood Estimation (MLE), we infer parameters quantifying remethylation kinetics of nascent DNA post-replication to individual CpG-site resolution, genome wide. Second, we perform stochastic simulation of various candidate enzyme-kinetic models of maintenance methylation in order to identify potential mechanisms consistent with the experimentally-inferred parameter distributions. Our combined statistical/mathematical modeling approach expands the utility of genomic datasets such as those resulting from Repli-BS experiments. The approach enables a basepair-level view of the combined influences of temporal and cell-to-cell heterogeneity across the genome.

## Methods

### Methods Overview

The workflow of our approach is summarized as follows. We analyzed published Repli-BS data [29], which tracks re-establishment of genomic methylation patterns in newly replicated DNA over time. A schematic of DNA remethylation process is shown in Fig. 1 A. We first employed analytical, stochastic models of remethylation kinetics to serve as a framework for analysis of the experimental data. These analytical models with few parameters (two to three) served primarily as a tool to quantify kinetics via statistical inference of post-replication DNA methylation, to single CpG-site resolution, genome-wide (Fig. 1 B and C). We then developed a set of candidate kinetic models of enzyme-mediated maintenance methylation; the aim in studying these more detailed and biologically motivated models was to provide mechanistic insight on maintenance methylation processes in conjunction with the inferred parameters from Repli-BS (Fig. 1 D). The connection between the two modeling frameworks (i.e., between the small analytical models with inferred parameters, and the more complex, enzyme-kinetic mechanistic models) was achieved as follows. We first present the primary outputs of the statistical inference: namely, (1) the distributions of per-site inferred parameter values across different chromosomes, (2) the correlation of parameter values with genomic distance (GD), and (3) the distribution of inferred parameter values with respect to local CpG density (CpG_*d*_). Next, we perform stochastic simulations of candidate enzyme-kinetic models, using parameters derived from previous literature where possible. Finally, we extract *in silico* Repli-BS read-data from the simulations, subject to the same experimental constraints (i.e., measured timepoints, read-depth) as the experimental data. We then compare outputs (1-3) from simulated and experimental read-data in order to assess the ability of different model mechanisms to reproduce features of the experimental outputs.

**Fig 1.**
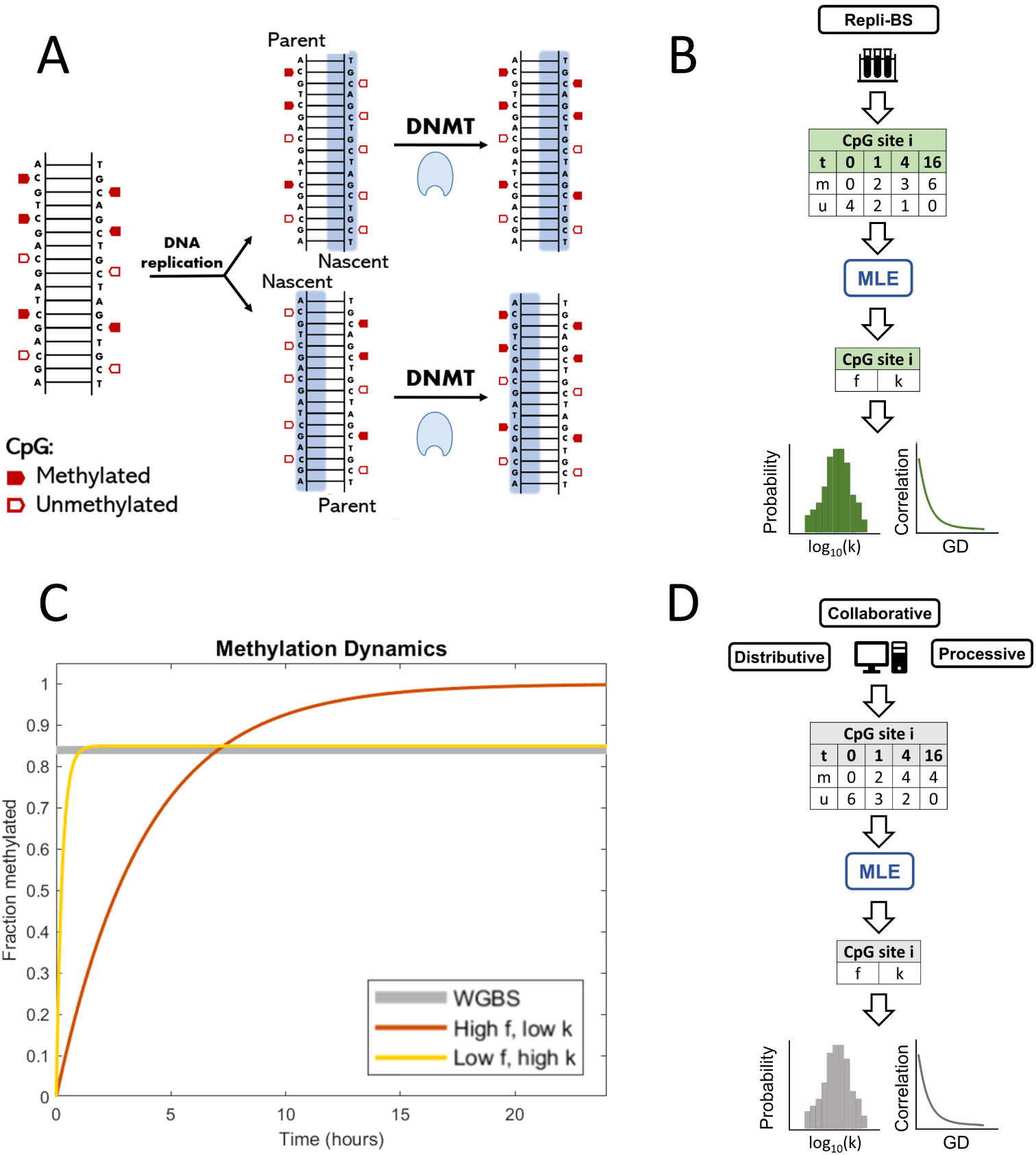
**A:** DNA methylation in the context of replication. Upon replication, complementary unmethylated nascent strands are synthesized for each parent strand, such that fully methylated CpGs become hemimethylated. Classically, full methylation is restored by DNMT1 (though DNMT3s have also been shown to contribute to this maintenance). **B**: Work-flow of the data analysis: Repli-BS data records methylated (m) and unmethylated (u) reads on the nascent strand for each site *i* genome-wide, with timpoints over 16 hours. MLE allows the inference of stochastic model parameters for each site *i*, giving outputs of parameter distributions and parameter-correlation with genomic distance (GD). **C**: The MLE procedure assesses remethylation rate (*k*) and steady-state remethylation fraction (*f*) for each CpG site, thus distinguishing between sites that are remethylated faster while reaching a lower average methylation level (yellow line) and sites with slower kinetics but higher methylation overall (orange line). In contrast, traditional WGBS would not distinguish these cases, as they have roughly the same time-average (grey line). **D**: Work-flow of the enzyme-kinetic simulations: Stochastic modeling of remethylation kinetics according to either a Distributive, Processive, or Collaborative mechanism generates simulated datasets, which are then analyzed with the same MLE procedure used for the experimental data, shedding light on *in vivo* mechanisms.

**Fig 2.**
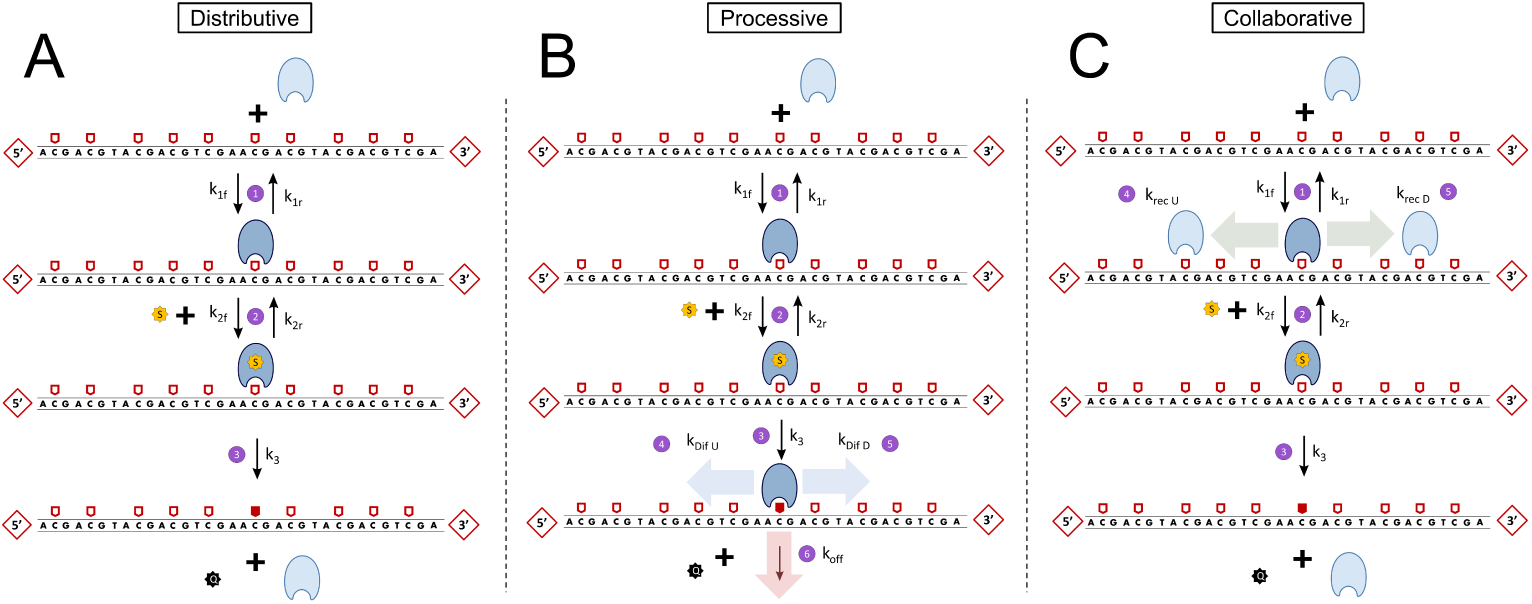
DNA remethylation Distributive (A), Processive (B) and Collaborative mechanisms (C). Hemimethylated sites (e.g. sites that can be remethylated) are indicated as empty pentagons, while methylated sites are represented as red-filled pentagons. Unmethylated sites are not represented in the scheme. In the Distributive model DNMT1 binds to a hemimethylated CpG site, incorporates SAM, and catalyzes methylation. Methylation and release of both DNMT1 and SAH occur in a single step. In the Processive model, after methylation DNMT1 can diffuse towards its immediate neighbor sites either upstream (U) or downstream (D). In the Collaborative model, once DNMT1 is bound to a hemimethylated CpG, a second DNMT1 molecule can be recruited onto nearby CpG sites. A distance function adapted from [42] favors recruitment at nearer CpGs.

### Experimental Data from Repli-BS

In the Repli-BS experiments [29], human embryonic stem cells (HUES64) were pulsed for one hour with bromodeoxyuridine (BrdU), and bisulfite sequencing measurements were obtained at multiple timepoints between 0 and 16 hr post-pulse. Methylation was measured on BrdU-labeled DNA, thereby selecting only those cells in which DNA replication occurred during the pulse interval ([-1,0] hours). The captured bisulfite read-data measured the presence (1) or absence (0) of methylation at individual CpG sites. Thus, the experimental data is of the form 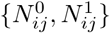, or observed numbers *N* of unmethylated reads (“0”) and methylated reads (“1”) on nascent DNA at each timepoint *j* at site *i*. Each measured site comprised a variable number of acquired reads at each timepoint. For parameter inference, we analyzed four timepoints (0, 1, 4, and 16 hr). In the experiments, nascent DNA (0 hr) was collected and analyzed from cells sorted according to their stage in S-phase of the cell cycle (stages S1-S6). We filtered methylation data from the 0 hr time point so as to retain only values from CpGs being actively replicated within each S-phase fraction, based on sequencing read enrichment (replication-timing). The remaining methylation values were then merged into a single 0-hr dataset for the first timepoint for our analysis. We restricted analysis only to those sites that had a minimum total read-depth of 15, with at least ten at time 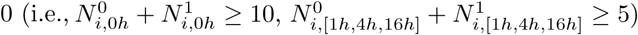. After these restrictions, the dataset contained 10,435,822 analyzed unique CpG sites, which constitute ≈ 40% of the total number of sites in the original set.

### Statistical Inference of CpG Remethylation Kinetics

#### Analytical kinetic models

We employed analytical, stochastic models of remethylation kinetics to serve as a framework for analysis of the experimental Repli-BS data. (Analytical means here that models admit a simple, analytical formula for the likelihood function used for parameter estimation). The basic model assumes that fully methylated CpG sites (i.e., dual-methylated on both strands) become hemi-methylated at the time of DNA replication, followed by subsequent remethylation of the nascent strand over time (Fig. 1 A). Each CpG site *i* is characterized by two parameters: *f*_*i*_ (the fraction of cells in the population that are methylated at site *i* in the steady-state) and *k*_*i*_ (the rate of remethylation at site *i*). Methylation of an individual, hemimethylated site is assumed to be an independent, memoryless, stochastic process. Under these assumptions, a CpG site in an individual cell, which is hemimethylated at the time of replication, has probability to experience remethylation at time *t* post-replication given by the exponential distribution:

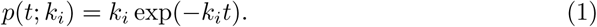

Thus, for a population of cells, the probability of observing a methylated read (denoted ‘1’) on the nascent strand at site *i* is given by

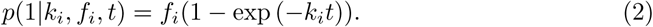

Since each read is 0 or 1, the probability of observing an unmethylated read (‘0’) is given by

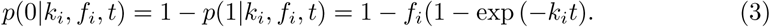

The half-life to remain unmethylated, or the “maintenance methylation lag-time” of an individual site *i* is then

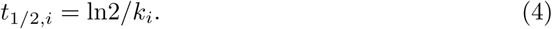

We emphasize that the primary utility of this simple model is to enable estimation of rate constants (and thus timescales) of remethylation kinetics across the genome from Repli-BS data. The inference results for this simple model with independent CpGs can nevertheless reveal more complex mechanisms of maintenance methylation, as described below (see Results).

The basic, two-parameter ({*k*_*i*_, *f*_*i*_}) model assumes that post-replication methylation is strictly irreversible, i.e., it enables description of only monotonically increasing methylation-fractions at any given site over time, and cannot account for any loss of methylation within one cell cycle. (Tje model can, however, account for passive demethylation, i.e. if *k* is too slow for parent-strand methylation levels to be fully re-established within one cell cycle). In light of the demethylating activity of TET enzymes, we also performed inference using a three-parameter, reversible model, with time-dependent probability of methylation given by:

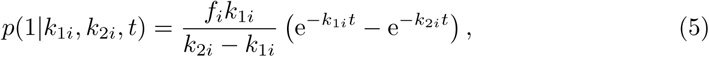

where *k*_1*i*_ is the rate constant to acquire methylation over time post-replication and *k*_2*i*_ is the rate constant of loss of methylation. This model provides the simplest means of fitting non-monotonic, reversible kinetics, potentially representing both methylation and active demethylation processes at a given site.

Due to the simplistic nature of the above models (eqns. 2 and 5), they can be considered to be agnostic to underlying biological mechanisms and to serve merely as tools to fit either irreversible or reversible kinetics. However, we can also draw a parallel between these models and other methylation dynamic model frameworks, which often comprise at least three types of sites, denoted *u* (unmethylated), *h* (hemi-methylated), and *m* (fully methylated). Equation 2 then accounts only for transitions *h* → *m*, and corresponds to a mean-field deterministic model where

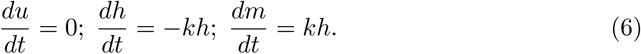

In contrast, the reversible model of Eq. 5 is in line with a model *h* → *m* → *u*, and mean-field equations

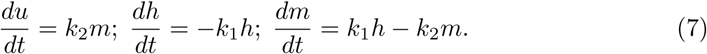

We stress that these equations, including in the reversible model case, neglect many biologically plausible reactions that are present in more detailed methylation dynamics models (e.g., *m* → *h*, etc.) in favor of enabling parameter fits to Repli-BS data for sub-cell-cycle timescales.

Both the two- and three-parameter models can be extended to incorporate sources of experimental error. Given a false-positive rate *E*_*p*_ (the probability of a false methylation count) and false-negative rate *E*_*n*_ (the probability of a false non-methylation count), then the probability of experimental observation of a methylation read is:

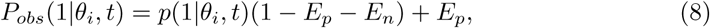

where the parameter vector *θ*_*i*_ = {*k*_*i*_, *f*_*i*_} for the two-parameter (irreversible) model and *θ*_*i*_ = {*k*_1*i*_, *k*_2*i*_, *f*_*i*_} for the three-parameter (reversible) model. The probability of experimental observation of an unmethylated read is *P*_*obs*_(0|*θ*_*i*_, *t*) = 1 − *P*_*obs*_(1|*θ*_*i*_, *t*).

In the above models, the time variable *t* denotes the time that has elapsed post-replication. More specifically, it can be taken as the instant at which nucleotides (including the thymidine analog BrdU) were added to the nascent DNA strand at a given locus. The experiments have inherent uncertainty related to this timing. Since the BrdU pulse was one hour in duration, replication could have occurred anytime within the hour-long pulse. As such, we convert the experimental “post-pulse” time to the model’s “post-replication” time in an unbiased way by adding one half hour. In other words, the experimental timepoint of 0-hour-post-pulse is converted to *t* = 0.5 hours post-replication (and similar for the other experimental timepoints). This conversion does not correct for any additional variability in replication timing that occurs within the pulse window. An alternative method, treating “time-post-replication” as a uniformly distributed random variable over the interval of one hour, was also studied (See Supplement).

#### Maxmimum Likelihood Estimation of Remethylation Rates

In order to estimate the model parameters (i.e., the rate and steady-state fraction-methylated), we ask how likely it is that the model would “produce” the measured experimental data. In general, for a model that describes the probability, *p*(**x**|*θ*) of an outcome given parameter(s) *θ*, the likelihood function is given by

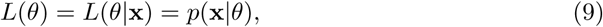

where **x** is the set of observations/outcomes. For *N* independent observations, the likelihood is thus

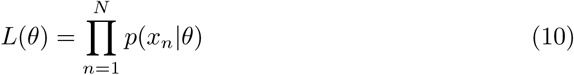

and the Maximum Likelihood Estimate of the parameters, given the data, is

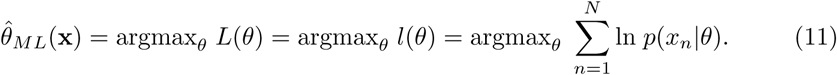

where *l*(*θ*) is the log-likelihood. The experimental data is of the form 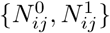, that is, observed numbers *N* of unmethylated reads (“0”) and methylated reads (“1”) at timepoint *j* at site *i*. Applying maximum likelihood estimation to the stochastic model of remethylation with parameters *θ*_*i*_ = [*k*_*i*_, *f*_*i*_] for site *i*, one obtains

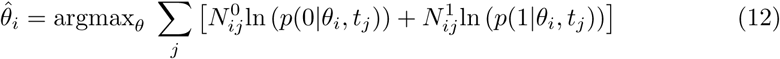

where *i* is the site index and *j* is the timepoint index.

In order to estimate the parameters, the log-likelihood surface *l* (*k, f*) was computed numerically for each set of site-specific read-data as a function of discrete *k* and *f* values, with domains *k* ∈ [10^*-*2^, 10], *f* ∈ [0, 1]. The limits in *k* were chosen by performing MLE for simulated data with the same timepoints and average read-depths as the experimental data, and identifying the approximate range of values over which *k*-estimation was possible.

The values of *k* and *f* for which the log-likelihood was maximum were taken as the estimated best-fit parameters for a given site. The exception to this was when the *k* maximum was located on the edge of the *k*-domain precluding unambiguous assignment (this generally only occurred on the upper edge, i.e., for very fast rates). In such cases, a Confidence Interval (CI)-based estimate of the lower bound on *k* was used (see below). All codes were written in MATLAB and will be available on Github.

#### Confidence Interval Estimation and Parameter Bounds

Confidence Intervals around ML estimates of the parameters for each site were computed using a Profile Likelihood method [47]. The Profile Likelihood corresponding to a specific value of a given model parameter, *σ*_*i*_ ∈ *θ* (where *i* here indexes the set of model parameters) refers to the maximum likelihood obtained when that parameter value is fixed while all remaining model parameters are freely varied. That is,

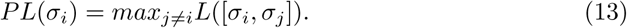

CIs for the parameter *σ*_*i*_ are then estimated by the range of values 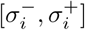 for which the Profile Likelihood falls within a defined range of the Maximum Likelihood,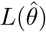. To approximate the 95% CI,

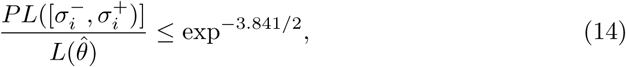

where the value of 3.841 derives from the 95th-percentile of a 1-degree-of-freedom *χ*^2^ distribution [48]. The estimation of ML parameters and Profile-Likelihood-based confidence intervals is represented in Fig 3.

**Fig 3.**
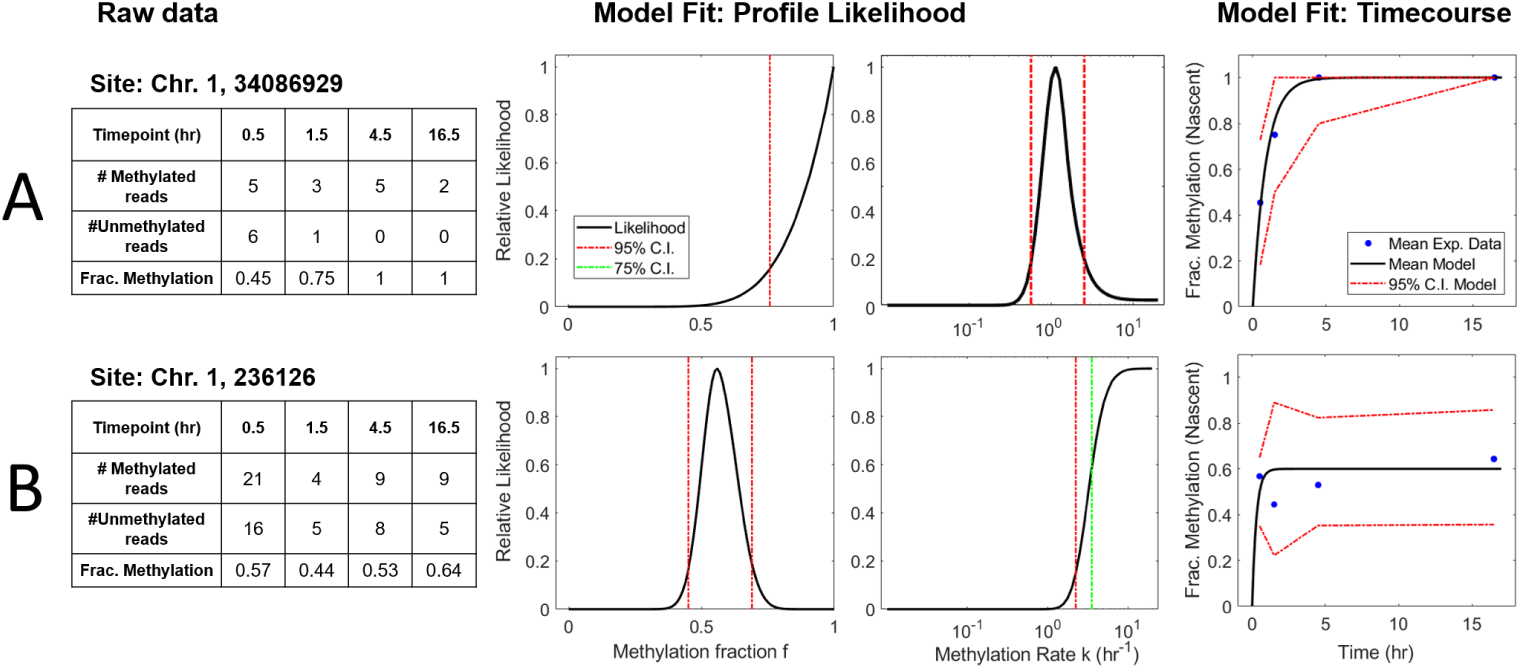
Representative read-data and model fits for two individual CpG sites on Chromosome 1. (A: site 34086929; B: site 236126) (left to right: raw experimental data, profile likelihood function for parameter *k*, profile likelihood function for parameter *f*, and model-predicted mean timecourse overlaid with experimental datapoints). **A:** Representative site with a global maximum in *k*, corresponding to parameter values *k* = 1.1 hr^*-*1^ ([0.56, 2.5] for the 95% CI) and *f* = 1 ([0.76, 1] for the 95% CI). The maximum-likelihood model prediction of mean fraction-methylation versus time (right, black curve) is overlaid with averaged experimental data (right, blue dots). The 95% confidence intervals for the model-predicted timecourse are simulated by Eqn. 2, while accounting for the variable number of samples (reads) at each timepoint. **B:** A representative site where the remethylation kinetic are too fast to measure, given the time resolution of experiments. In this case, the likelihood function increases to a plateau that extends infinitely in the direction of increased *k*, and only a lower bound on *k* can be determined unambiguously. In such cases, we take the lower 75% confidence bound as the parameter estimate for subsequent analysis (see Methods). Thus, the estimated parameter values for this site are *k* = 3.5 hr^*-*1^ ([2.2, ∞]) and *f* = 0.6 ([0.45, 0.69]).

#### Parameter Correlation Function

Correlation of inferred parameters is calculated as a function of genomic distance (i.e., number of basepairs separating individual CpG sites). As analyzed CpGs are unevenly spaced along the genome, correlation is calculated for binned distances [49]. The correlation function of parameter *θ* is given by

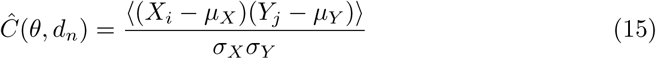

where *d*_*n*_ is the *nth* discrete distance bin, and (**X, Y**) are the pairs of parameters (*θ*_*i*_, *θ*_*j*_) = (*k*_*i*_, *k*_*j*_) or (*f*_*i*_, *f*_*j*_) at sites with genomic positions *i* and *j* where *d*_*n-*1_ *<* |*i* −*j*| ≤ *d*_*n*_. *µ*_*X*_ and *σ*_*X*_ are the mean and standard deviation, respectively, of the parameter values in **X** (and similar for **Y**). This definition is identical to that used in other analyses of correlated methylation fractions [43].

### Single-basepair-level stochastic enzyme-kinetic models

We performed simulations of single-CpG stochastic enzyme-kinetic models according to a set of candidate mechanisms, called the Distributive, Processive, and Collaborative models. These models focus only on the process of maintenance methylation, i.e., the remethylation process occurring over *<* 20 hours, and neglecting additional processes such as methyl erasure. In the three models, DNA is treated as a one-dimensional system of CpG sites which can be either unmethylated (u), hemimethylated (h), or methylated (m). Immediately after replication, sites are assumed to be in either the unmethylated or hemimethylated states, with hemimethylated sites being susceptible to remethylation by the enzyme (E, assumed namely to be DNMT1). The reaction also requires the methyl donor substrate, S, which stands for S-adenosylmethionine (SAM), while Q stands for its unmethylated form, S-adenosyl homocysteine (SAH). While sharing a common backbone in terms of E and S binding, as well as the remethylation reaction, the three models differ in the manner in which the enzyme reaches new hemimethylated sites after catalyzing methylation at one site.

#### Distributive mechanism

The backbone Distributive mechanism is based on a Compulsory-Order Ternary-Complex Mechanism (COTCM), by which DNMT1 (*E*) first binds the hemimethylated CpG (*h*) to form the *Eh* complex, and subsequently a SAM molecule (*S*) forming the ternary complex *EhS* (Fig. 2 A). While *m* stands for the methylated CpG, *Q* stands for SAH, the unmethylated form of SAM. The Distributive mechanism treats individual CpG sites as fully independent. The value of the forward and reverse rate constants for the first two binding reactions 1 and 2 (*k*_1*f*_, *k*_1*r*_, *k*_2*f*_, and *k*_2*r*_), as well as the catalytic step 3 (*k*_3_) have been derived from experimental values in [17] (See Supplementary Information for more details). (Note that there is no direct relationship between the kinetic constants here and in the analytical kinetic models). All parameter values for the this and the other models can be found in Supplementary Table S1.

#### Processive mechanism

The Processive mechanism assumes that DNMT1 can remain bound to DNA after performing the catalytic step and reach subsequent hemimethylated CpG sites by diffusing along DNA. The first two steps are identical to the Distributive model, with *E* binding *h* to form *Eh*, and *Eh* subsequently bonding *S* to form *EhS* (Fig. 2 B). The same assumptions were made to determine the rate constants associated with these steps.

In the Processive model, however, the catalytic step (*k*_3_) does not directly imply DNMT1 to drop off from the DNA chain, returning as a free species into solution. Instead, *E* remains bound to the DNA molecule onto the recently methylated site, forming the *Em* complex, while releasing a molecule of *Q*. Then, two different events can take place: on one hand, DNMT1 can move towards its neighbor CpG sites either upstream (towards the 5’ end) or downstream (towards the 3’ end) through linear diffusion along DNA. A new *Eh* complex with the destination site will be formed. Alternatively, DNMT1 can drop off the DNA chain and return into solution, with a rate constant *k*_*off*_. In both events, a methylated CpG site is left behind. The model is processive in the sense that DNMT1 can with high likelihood perform successive methylation on sufficiently close *h* sites. However, there is no explicit requirement in the model that DNMT1 move unidirectionally.

To incorporate diffusion into the stochastic simulations, we use a First Passage Time Kinetic Monte Carlo algorithm, based on ref. [50]. We assume the enzyme travels with 1D diffusion coefficient *D* and unbinds with rate *k*_*off*_. The algorithm requires computation of the probability that an enzyme will reach each of three “exit-states”: the nearest upstream neighbor *h* at distance *d*_*U*_, the nearest downstream neighbor *h* at distance *d*_*D*_, or the solution (by unbinding). The algorithm also requires computation of the First Passage Time density function, i.e. the distribution of waiting times at which the enzyme will first reach one of these three exit states, which is performed using Gillespie’s Eigenvalue approach [51]. Details of the Processive Model in can be found in the Supplement.

#### Collaborative mechanism

The Collaborative mechanism shares reactions 1,2 and 3 (and associated parameters) with the other two models. In this case, the catalytic step *k*_3_ implies enzyme drop-off after methylation, just like in the Distributive model. However, here sites are interdependent through a phenomenological mechanism of “collaboration” between enzyme molecules: after DNMT1 binds, a second enzyme can be recruited to any nearby CpG site upstream or downstream the original site (not necessarily the contiguous). The stochastic propensity for each recruitment reaction *k*_*RecR*_, notwithstanding, is indeed a function of the distance of a neighboring hemimethylated site to the recruiting site according to:

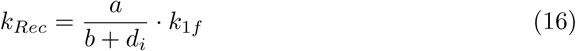

Where *d*_*i*_ is the distance between the recruiting and the neighbor hemimethylated CpG sites, and *a* and *b* are free parameters integrated into a non-dimensional distance-dependent function. Therefore, the recruitment propensity decreases with distance. The phenomenological model and the mathematical form of the distance function was adopted from a previous modeling study [52].

Note that classical views of a collaborative mechanisms are based on the fact that DNMT1 is recruited by agents such as UHRF1 [53]. Our model does not implicitly consider UHRF1, but through the distance-dependent function assumes that DNMT1 recruitment after a first copy is bound to DNA indirectly account for the fact after recruiting the first enzyme copy, UHRF1 can recruit a second copy close to it, hence being a simplification of a more complex biological reality.

#### Stochastic simulations

Stochastic simulation was carried out using the Stochastic Simulation Algorithm [54], except in the case of the Processive model as described above.

Before any reaction could take place, the first step consisted of simulating the substrate, i.e. DNA containing *N*_*sites*_ CpG sites, each assigned to be either unmethylated or hemimethylated at time 0 post-replication. Site numbers (and resulting inter-CpG distances) and methylation assignments were mined from an independent experimental dataset (WGBS measurements from Chr1 in arrested HUES64 cells, [29]) in the following manner: A start site was randomly sampled from Chr1, and the following contiguous *N*_*sites*_ measured sites from the WGBS dataset were taken as the population-average, steady-state methylation landscape for the simulated substrate (with *N*_*sites*_=100,000). Each site in a simulated “cell” was assigned a steady-state methylation status of m or u (i.e., methylated or unmethylated) randomly, with probability of methylation matching the population average; these assigned steady-state fractions are denoted *f*_*a*_. If a given site in a cell is assigned to be methylated at steady-state, then it is assumed to be hemimethylated at time 0, and kinetics of re-methylation proceed according to the model mechanisms described above. Unmethylated sites remain as such for the duration of the simulation. Note that data from arrested cells (which are not undergoing DNA replication) are chosen to estimate *f*_*a*_, as measurements in these cells are assumed to reflect a steady-state methylation landscape in the absence of replication-associated temporal dynamics.

Simulations of time-trajectories were performed for multiple cells in order to obtain multiple “reads” of methylation status for each site, in accordance with the experimental read-depth afforded by the Repli-BS data. In this way, simulated datasets were produced for each model *in silico* of the form 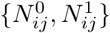 matching the experimental dataset in timepoints and distributions of read-depth for each site (see Supplement for details).

## Results

### Identification of Single-CpG Remethylation Kinetic Parameters

Statistical analysis of Repli-BS data by Maximum Likelihood Estimation enabled per-site inference of the rate of post-replication methylation of the nascent strand (*k*) and the steady-state fraction of cells methylated on the parent strand (*f*), according to Eqn.1. Note that the parameter *f* here has a slightly different meaning than the methylation fractions obtained traditionally from bulk WGBS data. As illustrated in Fig. 1C, WGBS typically averages methylation from asynchronously dividing cells, and thus captures DNA strands in different stages of time after replication. In contrast, *f* as inferred here represents the fraction methylation in the steady-state (rather than the time-averaged fraction methylation), that is, after the methylation status of a given site has been returned to the “baseline” delineated by the parent strand before replication.

The parameters were generally “identifiable”, that is, a single global maximum was present in the computed bivariate (*k* vs. *f*) likelihood surface, and the parameter values corresponding to this peak were thus obtained as the Maximum Likelihood Estimates. However, due to the limited time-resolution of the experiments, some sites experienced remethylation too quickly to allow unambiguous assignment of *k*. This occurred when all or nearly all reads were methylated from the earliest timepoint (0 hour post-pulse, estimated to be an average of 0.5 hour post-replication, see Methods). In the statistical analysis, this manifested as a one-sided plateau in the profile likelihood function of *k* (Fig.3 B), enabling only identification of a lower bound on the rate constant *k*. In such cases, we used the value of the lower 75% Confidence Interval as an estimate of *k*, reasoning that this provides a conservative estimate of the remethylation rate given the experimental time resolution. Note that the parameter *f* is by definition bounded, *f* ∈ [0, 1], so the maximum likelihood value of *f* frequently occurred on the edge of the domain (Fig.3 A). In such cases the profile likelihood function increased steeply toward the edge (i.e., toward *f* = 1), and since *f* is also by definition bounded, we directly used the edge-located maximum in *f*, rather than using CI-determined bounds as with *k*.

The distributions of inferred parameters for Chromosome 1 according to the two-parameter model (Eq. 2) are presented in Fig. 4. The distribution of *f* values shows a bimodal pattern, similar to the directly-measured fractions from WGBS (*WGBS*_*f*_), with most sites having high methylation fractions (*f* > 0.6, with the majority showing full methylation, *f* = 1) and a small subset of unmethylated sites (*f* =0). The distribution of inferred *f*-values is qualitatively consistent with previous WGBS measurements in cell-cycle-arrested cells (see Fig. S9), though it shows a relatively smaller fraction of *f* = 0 sites. Our read-depth restrictions were more likely to exclude these unmethylated CpGs, likely resulting from asymmetric PCR amplification efficiencies of methylated versus unmethylated strands [55]. Overall, the results support that our inferred *f* fractions here can be considered to be analogous to WGBS-derived fractions, albeit “corrected”, i.e., with the influence of replication-associated kinetics removed.

**Fig 4.**
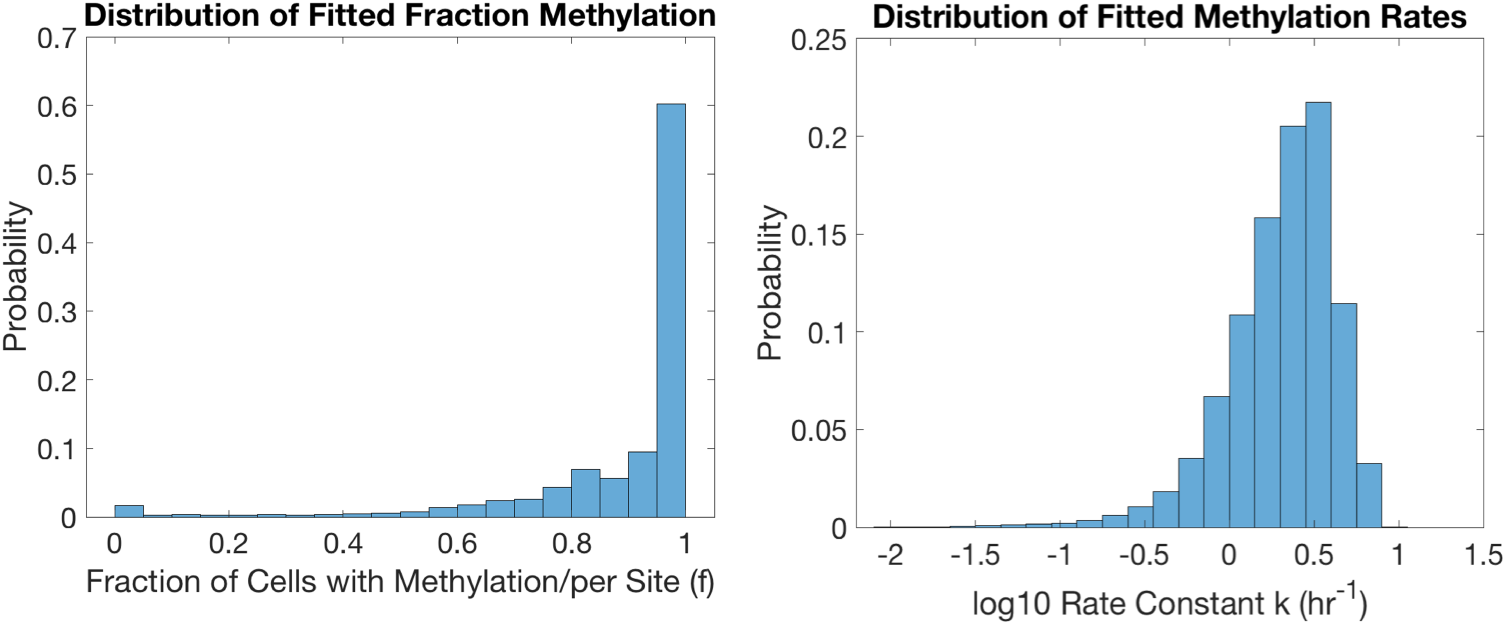
Histogram of inferred steady-state fraction methylation, *f* (left), and remethylation rate, *k* (right) for Chromosome 1 in hESCs. Histograms represent ∼ 0.8 million measured CpGs, and are normalized by probability.

The distribution of remethylation rate *k* values shows high site-to-site variability in remethylation kinetics. Non-zero *k* values were estimated over a range from 0.01 to 9.5 hr^*-*1^, with 95% of the values lying within 0.3 and 6 hr^*-*1^ ([2.5%,97.5%]). That is, 95% of sites were found to have a “half-time to remethylation” between > 2 hours and *<* 10 minutes. A small fraction of sites had *k* below 0.1 hr^*-*1^, or a half-life of 7 hours or more. The median value of inferred *k* for Chromosome 1 was 2.2 hr^*-*1^. As noted previously, for the fastest sites, the identified constant *k* can only be taken as a lower bound for the true rate; as such, the true *k* − distribution is likely wider than that presented in Fig. 4, and the curtailed shape of the *k* − distribution on the right-hand-side likely reflects the experimental limit in time-resolution, rather than the true kinetic distribution. Since kinetics of remethylation are meaningless for sites that remain unmethylated, only sites with *f* > 0 have an associated estimate for *k*.

The parameter estimates in Fig. 4 are based on a model (Eq. 2) that assumes monotonic remethylation kinetics, i.e., in the several hour time window post-replication, methylation on the nascent strand is assumed to increase or the site remains unmethylated, but it cannot decrease. To relax this assumption, we also estimated parameters for the 3-parameter “reversible” methylation kinetic model (see Methods, Eq. 5). Using a Bayesian Information Criterion-based model selection, we found that < 1% of analyzed CpG sites on Chromosome 1 were better fit by the reversible model (see SI), concluding that for the majority of sites, the monotonic 2-parameter model sufficiently captures the kinetics revealed by the Repli-BS measurements.

Error analysis of the parameter estimates was performed in various ways. We generated “ground truth” simulated data with identical timepoints and read-depth distribution as the experimental data, and then tested ability of the MLE approach to recover the correct parameters (See MLE Validation in SI for details). In this analysis, the broad features of *in silico*-assigned parameter distributions were recovered accurately (Fig. S2). The error in individual estimates of *k* ranged widely depending on the assigned values of *k* and *f* and the selected read-depth. We estimate an overall average relative error in per-site *k* values of approximately 32%. The average absolute error in inferred *f* values was 0.1. Overall, we concluded from this analysis that individual inferred parameters can be subject to relatively large error, while broader features of the distribution can be accurately inferred. Moreover, individual-site *k*’s can be estimated with high confidence to within an order of magnitude. For example, for assigned *k*^*I*^*s* of 1 hr^*-*1^ (in the mid-range of the inferred distribution), 50% of inferred values fell within 0.71 and 1.41 hr^*-*1^, and 95% of estimates fell within 0.50 and 2.51 hr^*-*1^, which is less than one order of magnitude (see Fig. S2).

We varied the method of parameter estimation to determine whether the parameter distributions shown in Fig. 4 are robust to details of the estimation method. First, we tested the influence of an experimentally-informed Bayesian prior on the estimated parameters. As discussed above, per-site methylation fractions obtained in the same cell line from WGBS measurements in arrested cells are expected to be a reasonable independent measurement of our statistically inferred values of *f*. We therefore used these independent measurements to construct Bayesian priors on *f*_*i*_, and determine the impact on the estimates *k*_*i*_. We found that some individual per-site estimates were affected by this choice of prior, but in general the bulk of estimates were similar between the two approaches and the broad characteristics of the distribution were unchanged (see Fig. S4). In another approach, we tested whether including explicit treatment of unknown sources of experimental measurement error in the model (Eq. 8) would affect the estimates of *k* and *f*, and found that within standard ranges of error estimates the distributions remained largely the same (see Fig. S5).

### Inferred parameters reveal high intra-chromosomal variability, but little variation between chromosomes

While the inferred parameters, remethylation rate *k* and fraction methylation *f*, show high variability from site to site, their distributions are highly uniform across different chromosomes. Representative *k* distributions are presented in Fig. 5. (see also Fig. S7 and S8).

**Fig 5.**
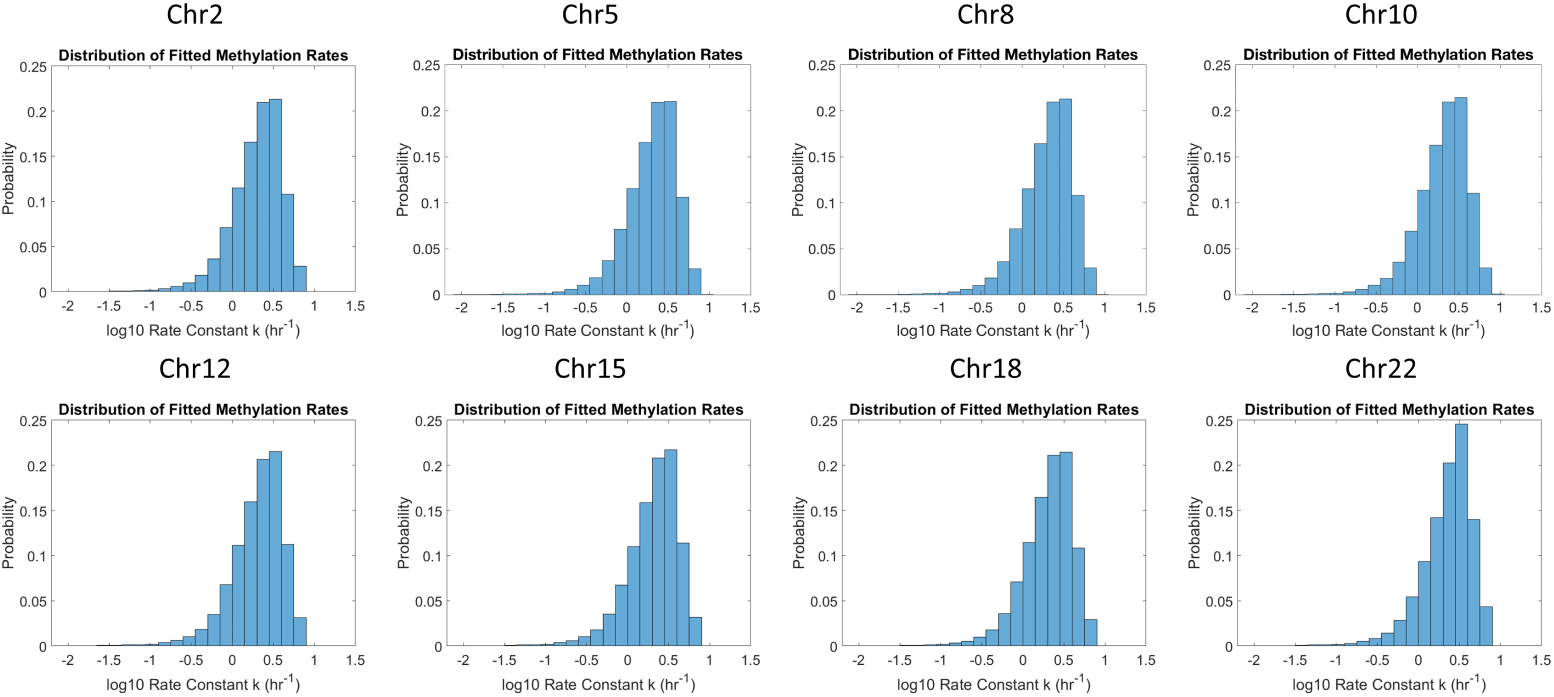
Histogram of remethylation rates, *k*, for Chromosomes 2, 5, 8, 10, 12, 15, 18, 22. Histograms are normalized by probability.

### Inferred parameters correlate with local CpG density

Individual-CpG-site inference of *k* and *f* allow the study of how both parameters depend on local CpG density (CpG_*d*_) (Methods, Eqn. X). In general, higher density regions show more CpGs with lower methylation fractions (Fig. 6). The mean value of *f* for low, medium, and high density regions was 0.91, 0.90, and 0.75, respectively. These averages reflect increasing probability of sites in each density group with *f* = 0. This result is in agreement to what other authors have reported for WGBS methylation fraction: high-CpG_*d*_ areas are more likely to be hypomethylated than low CpG_*d*_ areas [56]. The distributions of inferred *k* parameters also show dependence on CpG_*d*_. In general, the distributions shift rightward with higher density, i.e., faster remethylation is associated with higher CpG_*d*_. However, in the highest density group there is also the appearance of an extended tail toward lower rates. Together these features give rise to a nonmonotonic dependence of average rate on density; mean remethylation rates *k* were 2.4, 3.1, and 2.5 *hr*^*-*1^ in the low, medium, and high density groups, respectively.

**Fig 6.**
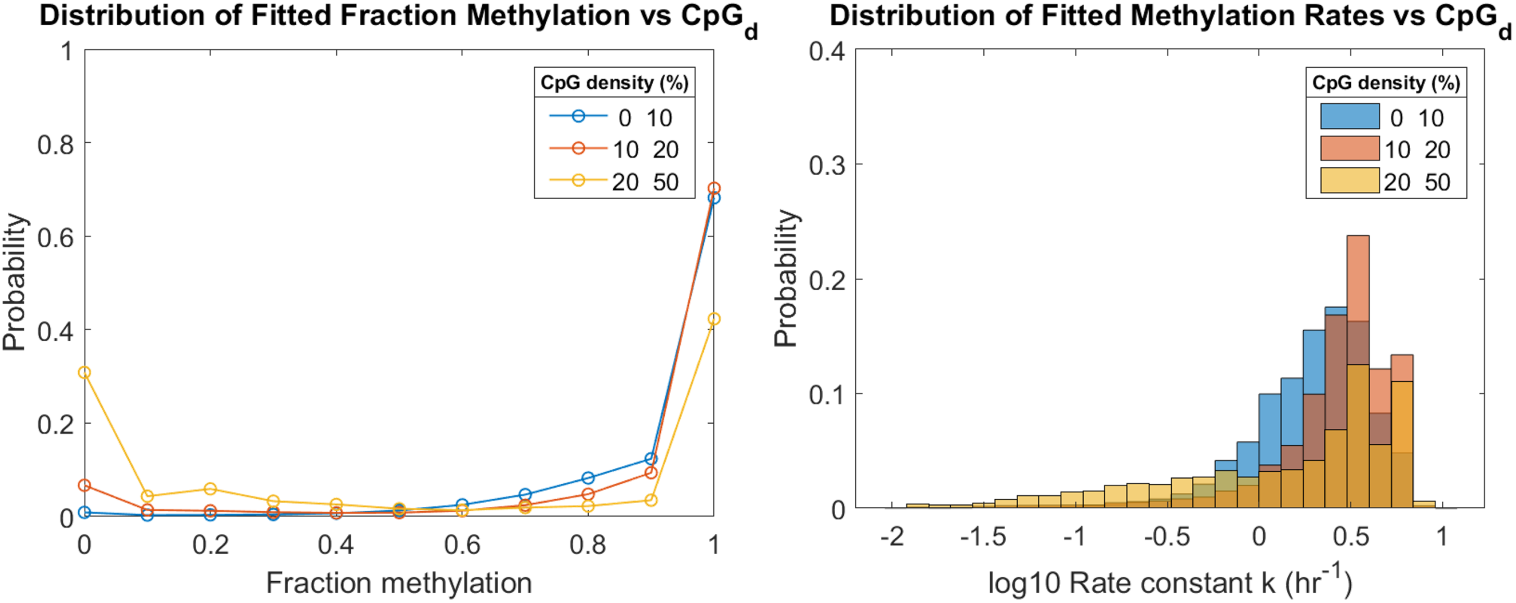
Remethylation rates and fraction methylation distributions for low, medium, and high-density CpG sites of Chr1. CpG_*d*_ of a site *i* is determined as the fraction of *bp* that are part of a CpG dinucleotide within a radius of 50 *bp* upstream and downstream the DNA molecule. Low-density CpG-sites represent 87% of the total sites analyzed in Chr1. Medium-density CpG sites represent a 12%, and high-density sites less than 1%. Low density is defined as [0,10)%. Medium density is defined as [10,20)%. High density is defined as 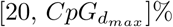, where 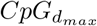 is the maximum *CpG*_*d*_ found in Chr1 (50 %).

Similar correlations between *k* and CpG_*d*_, as well as between *f* and CpG_*d*_, are observed along the genome (See Fig. S10 and S11 respectively).

### Remethylation parameters are correlated among neighboring sites

Individual-CpG-site estimation of kinetic parameters enables analysis of correlation between parameters of neighboring sites (see Methods, 15). We computed correlation as a function of genomic distance (GD), based on the individual CpG site IDs for analyzed sites (Fig. 7).

**Fig 7.**
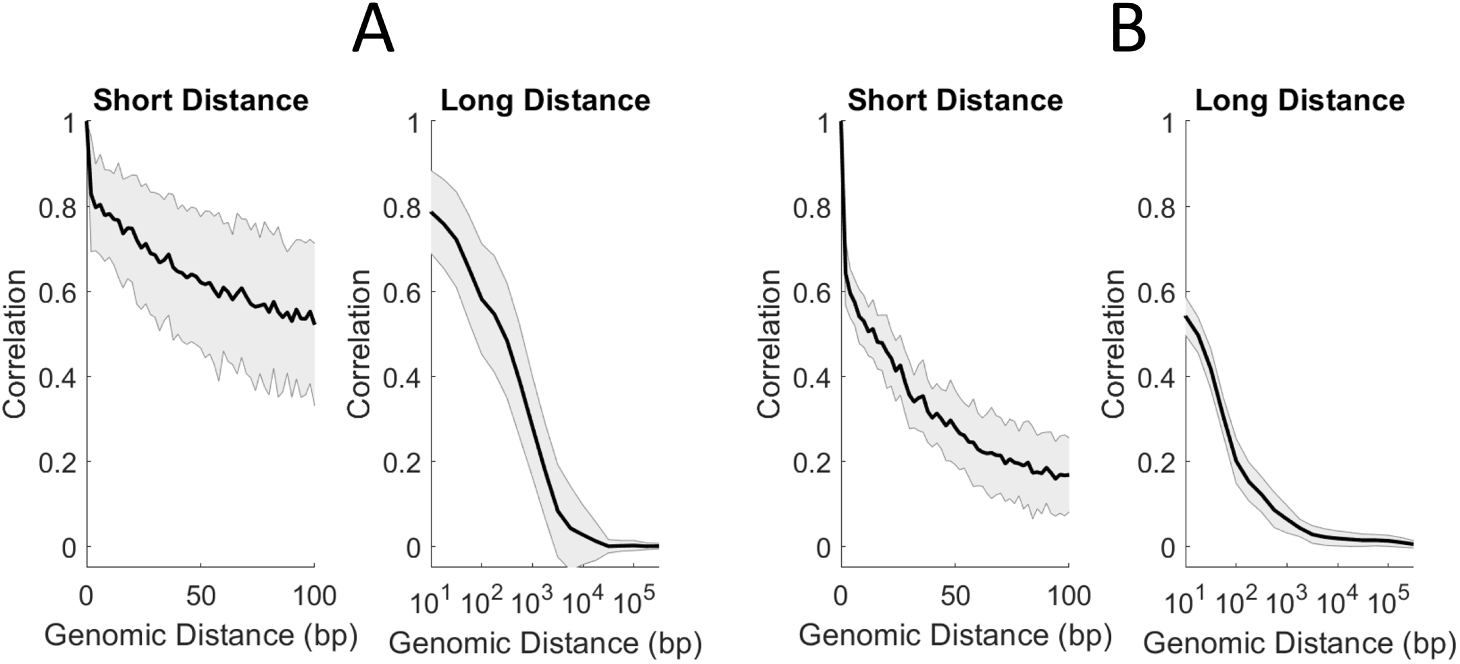
Correlation of fraction methylation *f* (A) and remethylation rates *k* (B) with GD. Correlation over short distances (left) and long distances (right) for Chr1.

We found that both parameters were correlated on neighbor sites, albeit with different shapes and lengths of their correlation functions. *f*-correlations are stronger than *k*-correlations for sites in the immediate vicinity: for example, adjacent CpGs on Chromosome 1 have an average *f*-correlation of 0.83 and an average *k*-correlation of 0.64. The *f*-correlation first drops below 0.5 at a distance of approximately 300 bp, while for *k* this dropoff to < 0.5 occurs around only 16 bp. Despite this more rapid initial dropoff in *k*-correlation, *k* values appear to have a weak but consistent correlation that persists out beyond 10 kilobasepairs. As with the distributions and density-dependence, the correlation functions showed remarkable uniformity across different chromosomes (Figs. S12 and S13).

### Different enzyme models produce distinct parameter correlations with CpG distance and density

In order to further understand the inference results from Repli-BS data and gain mechanistic insight into the process of maintenance methylation, we studied three mathematical models encoding different candidate mechanisms for DNMT1-mediated remethylation post-replication (Fig. 2). First, we employed a Distributive mechanism in which remethylation at each CpG site is independent from the surrounding sites. Second, we employed a Processive mechanism in which DNMT1 can linearly diffuse along DNA after methylating one site, potentially accessing contiguous hemimethylated neighbor sites in this manner. Finally, we employed a Collaborative mechanism in which DNMT1 can be recruited onto nearby hemimethylated CpG sites after a first enzyme-copy is bound on a nearby site. These different mechanisms capture aspects of previous mathematical models of maintenance methylation (see Methods).

For each model, we performed stochastic simulations of the remethylation process over a 17 hour period post-replication in order to generate simulated read-data with the same characteristics as the experimental Repli-BS data. Simulations were performed for DNA substrates containing 100, 000 CpG-sites (comprising ≈4.5 million bp), with steady-state methylation landscapes derived from experimental data (see Methods). We then analyzed the per-site kinetics of the simulated data with the same MLE procedure as used for the experimental data. In this way, we could determine the effect of the more complex enzyme-kinetic mechanisms on the per-site inferred kinetics. We could furthermore determine which model mechanisms generated data features in common with the Repli-BS experiments.

When using the different molecular mechanisms to stochastically simulate remethylation kinetics, using parameter derived from enzyme kinetic studies [17], we observed distributions in per-site *k* parameters (*k*_*model*_) that are somewhat slower overall and narrower than the corresponding experimental distributions (Fig. 8, Top). Furthermore, the *k*_*model*_ distributions are generally similar for the three models.

**Fig 8.**
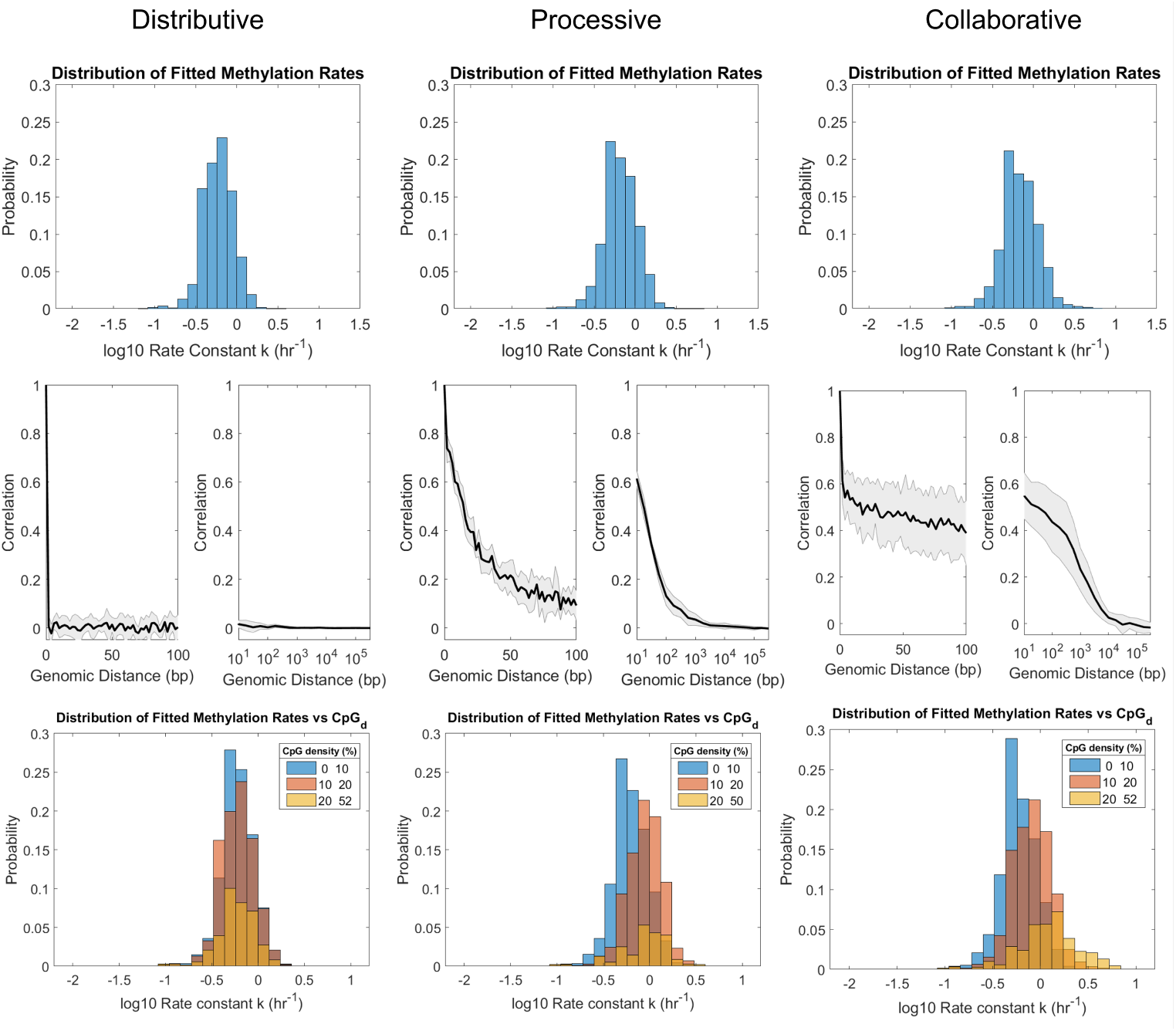
Simulated remethylation rates histograms (Top), *k*-correlation with GD (Middle), and *k*-dependence on local CpG_*d*_ (Bottom) when using each of the proposed mechanisms (Distributive, Processive, and Collaborative). The same *in silico* cluster containing 10^5^ sites was used for all models. Both the position and the *f*_*a*_ for every site where sampled from an independent experimental dataset of WGBS measurements from Chr1 in arrested HUES64 cells [29].

Major differences appear in terms of *k*_*model*_ correlations with GD (Fig. 8 Middle). Kinetic rates derived from the Distributive model do not correlate with GD to any extent, i.e., the correlation function immediately drops from 1 for *GD* = 0 (a given site is fully correlated with itself) to fluctuate around 0 for all *GD* ≥ 2 (the minimum distance between CpG dinucleotides). In contrast, *k*_*model*_ values derived from the Processive and the Collaborative mechanisms show distinctive correlation functions that decrease with GD. The precise shapes and persistence of the correlation functions of both Processive and Collaborative mechanisms were found to depend on the models’ parameters (Fig. S14 and S15), but the existence of correlation is robust. In contrast, the Distributive model cannot produce neighbor correlations for any choice of model parameters. Overall, these results show that correlation between kinetics on different CpG sites is not imposed by the local features of steady-state methylation patterns, nor by the MLE procedure itself (as these are common to all three models). Rather, the neighbor-correlations result from the DNMT1-mediated interdependence between neighboring sites. Moreover, the results show that neighbor correlation can result from two disparate mechanisms of inter-dependence (single enzyme processivity versus cooperation between multiple enzyme molecules).

Remethylation rates generated from the Processive and the Distributive mechanisms also show dependence on CpG_*d*_, with faster *k*_*model*_ values inferred for higher-density sites (Fig. 8 Bottom). This observation is in agreement with the mechanistic aspects of both models, since proximity between sites increase both diffusing and recruiting propensities. Again, the results of both models are in stark contrast with remethylation rates derived from the Distributive mechanism, for which *k*_*model*_ distributions remain centered around the same value for the three density groups. The shifted distributions of the Processive and Collaborative models with *CpGd* are in partial agreement with the experimental results of Fig. 6, where the bulk of the distribution is also seen to shift to higher *k* values with increasing density. However, none of the models capture the extended slow-kinetics tail observed experimentally for the high-density group seen in 6 and Fig. S10.

## Discussion and Conclusion

In this work, our approach combining statistical inference and mathematical modeling reveals genome-wide temporal heterogeneity in DNA methylation maintenance kinetics. Inferred kinetic rates of maintenance methylation varied by about two orders of magnitude across individual CpG sites in hESCs. The results further show how kinetics of maintenance methylation at individual CpGs depends on local CpG density and correlates with kinetics on neighboring sites. Stochastic simulations revealed that these correlations could be introduced by enzyme-mediated remethylation through either a Processive or Collaborative mechanism.

Our mathematical modeling-aided analysis approach helps to extend the utility of genomic datasets emerging from techniques such as Repli-BS to shed light on processes of epigenetic regulation. Specifically, the approach implemented here gives a deeper understanding of sources of IM, by disentangling heterogeneity in DNA methylation levels resulting from replication-associated temporal heterogeneity (i.e., due to lag in remethylation) versus stable cell-to-cell differences (Fig.1 C). In terms of the inferred parameters, sites with slower remethylation kinetics (lower *k* values) experience a longer delay in methylation inheritance and thus exhibit this type of temporal heterogeneity. While the biological significance of sites exhibiting this pronounced lag is not clear, Charlton and Downing, et al. suggest that hESCs show a more pronounced genome-wide lag while IM levels were reduced in more specialized cell types. A separate study reported that DNA methylation is relatively stable during replication in primary dermal fibroblasts [28]. Together, these observations suggest the lag may have a potential role in fate specification; for example, a delay in methylation restoration could provide transcription factors with a “window of opportunity” to bind methylation-protected loci [57]. Our results suggest that regulation of such a window is dynamic and temporally heterogeneous within a population of hESCs. Given that various forms of cellular heterogeneity are known to play key roles in stem cell fate decision-making and embryonic development [58], we posit that it will be important to develop a more comprehensive picture of how post-replication methylation timing and variability impact probabilistic differentiation systems.

Our results can potentially aid in the development of more detailed mathematical models of DNA methylation dynamics; for example, the results indicate that remethylation rates vary by two orders of magnitude or more across the genome. The observed broad distribution of kinetic rates may reflect the multiple ways in which DNMT1 can reach hemimethylated CpGs, i.e., directly and independently from solution, from neighboring sites (e.g., as in the processive and collaborative mechanisms), or through additional recruitment mechanisms that were not studied here. For instance, active recruitment of DNMT1 to the replication fork may account for the fastest inferred rates, as discussed previously [29]. The variations in kinetic rates could also be the result of increased competition between other DNA-associated factors and DNMT1 for CpG sites. In this case, variation in kinetic rates would reflect accessibility of hemimethylated DNA based on the local chromatin landscape. These types of recruitment/competition are not present in any of the models studied here, potentially explaining why the mathematical models (with *in vitro*-derived parameters) showed consistently slower and more narrowly-distributed kinetics as compared to the Repli-BS-inferred parameters.

For parameter inference, we chose simplistic analytical models (i.e., the two- and three-parameter models, Eqs. 2 and 5, respectively). The bulk of the analysis and modeling effort of this paper was then focused on the two-parameter model, since a model selection procedure indicated that the vast majority of sites were better described by this “irreversible” model. The rationale for applying such simplistic models for our initial inference was, first, that they impose minimal mechanistic assumptions on the observed data, and second, that the few-parameter models afforded straightforward application of MLE for parameter estimation. Our initial parameter inference of *k* and *f* assumes only irreversibility of post-replication methylation; it makes no assumption of which molecular species control the reactions or by what mechanism. Thus, this model cannot capture the full complexity of methylation dynamics (neglecting, e.g., active demethylation), but we employ it for its expedience in analyzing the Repli-BS results for the majority of measured CpGs.

Previous studies have used statistical inference to quantify per-site parameters governing maintenance methylation dynamics [35, 40]. A key difference between those studies and this one is that parameters in those and other models [33, 34, 36–38] quantified the probability of methylation to be correctly reestablished before the next round of division, whereas our parameters quantify per-hour kinetic rate constants on a sub-cell-cycle timescale. Therefore, the focus and scope of previous *in vivo* inference/modeling efforts has been on the stability of methylation patterns over longer timescales, e.g., over hundreds of generations [33] or over days to weeks in the context of epigenetic reprogramming [38], whereas the scope of our study is the enzyme-kinetic processes occurring within one round of replication. As such, one unique feature of our study is that it more closely connects the enzyme-kinetic literature on DNMTs with statistical analysis of genomic data. The temporal nature of Repli-BS experiments enables this connection. The difference in timescales between our model and others may also account for the suitability here of an irreversible model that neglects active demethylation: whereas the interplay of TET, DNMT3a/b, and DNMT1-mediated processes has been shown to be necessary to account for overall stability of epigenetic inheritance over many generations [41, 59], we found that the classical model of DNMT1-mediated maintenance methylation described reasonably well the reestablishment of methylation at most CpG sites within one cell cycle. Nevertheless, the ≈1% of sites at which reversible kinetics was apparent could possibly reveal loci of preferential active demethylation and be of future interest.

Interdependence of CpGs in methylation dynamics has also been the subject of previous modeling studies. Multiple mechanisms have been suggested for this interdependence [39–44, 59], from the enzymatic behavior of DNMT1 itself (e.g., through processivity) or in conjunction with other molecular species. This interdependence was found to play an important role in the stability of methylation inheritance [39, 42]. Processivity was suggested by biochemical studies, in which longer oligonucleotides experienced faster methylation kinetics [16, 46]. A linear diffusion model (which we consider a type of processivity because it often results in sequential methylation of neighboring sites) was previously found to be consistent with the enzymatic behavior of DNMT1 [39]. Additional phenomenological models of interdependence were introduced in [41, 42], one of which we adapted for use herein. We found that the presence (versus absence) of neighbor correlations was robust to other details of the models, such as the other kinetic constants and the initial conditions of the DNA methylation landscape. Therefore, the *k*_*model*_-GD correlation can be unequivocally attributed to enzyme-kinetic mechanisms of Processivity and/or Collaboration. This idea is reinforced in light of the fact that the three mechanisms share a common “backbone” in terms of the reactions they feature, with two sets of binding reactions and a catalytic step. In a similar way, *k*_*model*_ correlation with *CpG*_*d*_, is again only observed for the Processive and Collaborative models. Overall, these findings support the hypothesis that the rate of remethylation of one site is affected by the state and the position of surrounding ones, and show how independent-site-inferences can nevertheless reveal interdependence and thus reflect more complex mechanisms.

In future studies, it may be possible to use the specific features of the inferred correlation functions to shed light on the enzyme-mediated mechanism of remethylation *in vivo*. From our simulations, the linear diffusion model is more consistent with the rapid fall-off, but low-persistent *k*-correlation inferred from data, and as such appears more in line with experiments than the single Collaborative model studied here. However, we note that the specific features of the simulated correlation functions depend on a number of unknown parameters (see Supplement), and comprehensive parameter optimization for the enzyme kinetics is outside the scope of the present work. As such, we conclude both the Processive and Collaborative models to be broadly consistent with the Repli-BS data. However, neither of these models perfectly matched the experiment-derived correlation functions, nor did they account for the apparent bimodality of *k*’s in high density CpG regions. Therefore, our inference results may suggest more complex mechanisms of density/neighbor dependence in future studies.

A limitation of the current work is the various sources of uncertainty that contribute to individual site-estimates. A major source of uncertainty in the MLE estimates is the variable, often low, read-depth for individual sites. (We note that the individual site-uncertainties, for example, as quantified by the width of 95% confidence intervals from profile likelihood functions, depend not only on read-depth but also on the observed kinetics and their relationship to the available experimental timepoints.) Consideration of read-depth leads to a necessary tradeoff between the minimum per-site read-depth admitted for analysis and the total number of sites maintained in the analysis (here, 40% of the original set with the chosen restrictions). We sought to balance these factors and found through in silico “ground-truth” testing and alternative analysis methods that, while individual site estimates could be susceptible to significant uncertainty, the shapes of parameter distributions and their correlation, e.g., with GD, were robust. Additional sources of error include the hour-long BrdU pulse window that limits resolution of precise time-of-replication, the number and/or choice of experimental timepoints, and experimental errors in bisulfite conversion. Future experiments on sub-cell-cycle timescales with additional experimental conditions or increased sampling should enable an increasingly detailed understanding of maintenance methylation kinetics and, more broadly, of DNA methylation heterogeneity.

